# Phenotypic plasticity, but not genetic adaptation, underlies seasonal variation in the cold hardening response of *Drosophila melanogaster*

**DOI:** 10.1101/691741

**Authors:** Helen M. Stone, Priscilla A. Erickson, Alan O. Bergland

## Abstract

In temperate regions, an organism’s ability to rapidly adapt to seasonally varying environments is essential for its survival. In response to seasonal changes in selection pressure caused by variation in temperature, humidity, and food availability, some organisms exhibit plastic changes in phenotype. In other cases, seasonal variation in selection pressure can rapidly increase the frequency of genotypes that offer survival or reproductive advantages under the current conditions. Little is known about the relative influences of plastic and genetic changes in short lived organisms experiencing seasonal environmental fluctuations. Cold hardening is a seasonally relevant plastic response in which exposure to cool, but nonlethal, temperatures significantly increases the organism’s ability to later survive at freezing temperatures. In the present study, we demonstrate seasonal variation in cold hardening in *Drosophila melanogaster* and test the extent to which plasticity and adaptive tracking underlie that seasonal variation. We measured the cold hardening response of flies from outdoor mesocosms over the summer, fall, and winter. We bred outdoor mesocosm-caught flies for two generations in the lab and matched each outdoor cohort to an indoor control cohort of similar genetic background. We measured the cold hardening response of indoor and field-caught flies and their laboratory-reared F1 and F2 progeny to determine the roles of seasonal environmental plasticity, parental effects, and genetic changes on cold hardening. We also tested the relationship between cold hardening and other factors, including age, developmental density, food substrate, presence of antimicrobials, and supplementation with live yeast. We found strong plastic responses to a variety of field- and lab-based environmental effects, but no evidence of seasonally varying parental or genetic effects on cold hardening. We therefore conclude that seasonal variation in the cold hardening response results from environmental influences and not genetic changes.

## Introduction

All organisms residing in temperate climates must cope with seasonal fluctuations in their environment. Many species exhibit phenotypic plasticity, which grants them the flexibility to thrive during the growing season and survive unfavorable times. For example, aspects of cold tolerance are known to vary as a function of seasonal exposure and provide a mechanism for some species to successfully overwinter (Esterbauer and Grill 1978; Anderson *et al* 1992; Shearer *et al* 2016). While phenotypic variation can arise as a result of environmentally triggered plasticity, genetic variation in seasonally advantageous traits also exists (Dobzhansky and Ayala 1973; reviewed in Tauber and Tauber 1981 and Williams *et al* 2017). Therefore, genotypes that underlie variation in seasonally relevant phenotypes may change in frequency across seasonal timescales for short-lived organisms (King 1972; Grosberg 1988; Hazel 2002; Schmidt and Conde 2006; Behrman *et al* 2015). In the present study, we examine the relative importance of plasticity and rapid, seasonal adaptation in the cold tolerance of *Drosophila melanogaster*.

*D. melanogaster* is an ideal system for contrasting the importance of phenotypic plasticity and rapid adaptation as mechanisms for survival under seasonally fluctuating conditions. Notably, *D. melanogaster* has a short generation time, producing 10-15 generations per growing season (Pool 2015), and experiences dramatic changes in selection pressures across seasons that elicit rapid adaptation in life history traits (Behrman *et al* 2015). Populations of flies living in orchards evolve over the period of months (Bergland *et al* 2014) as they track changing fitness optima influenced by seasonal fluctuations in selection pressure (Machado *et al* 2018). Although many life history and stress tolerance traits have been shown to exhibit adaptive tracking across short timescales (e.g. Cogni *et al* 2014, 2015; Behrman *et al* 2015, 2018), some of these traits are also highly plastic (e.g. Chippindale *et al* 2004; Ayrinhac *et al* 2004). In general, when environmental pressures vary over timescales briefer than the lifespan of the organism, plasticity—as opposed to adaptive tracking—is more likely to occur (Levins 1968; Botero *et al* 2015). For instance, physiological responses to temperature may be more likely to exhibit plasticity because temperature can change rapidly over short timescales. In the present study, we examine the roles of plasticity and rapid adaptation in seasonally varying phenotypes using cold hardening in *D. melanogaster*.

Many species of insects plastically adapt to cold seasons via cold hardening, a phenomenon in which brief pre-exposure to cool temperatures results in greater cold tolerance (Chen *et al* 1987; Lee *et al* 1987; Lee *et al* 2006). *D. melanogaster* is capable of cold hardening, with an increase in cold tolerance resulting from pre-exposure periods as brief as half an hour (Czajka and Lee 1990). Cold hardening has been documented in larvae, pupae, and adults. (Rajamohan and Sinclair 2008; Kostal *et al* 2011; Jensen *et al* 2007). Cold hardening in flies is associated with widespread transcriptional changes (Qin *et al* 2005; MacMillan *et al* 2016), shifts in metabolite profiles (Overgaard *et al* 2007), and altered lipid composition of cellular membranes (Overgaard *et al* 2006; Lee *et al* 2006). Collectively, these responses permit maintenance of neuronal homeostasis under cold stress (Armstrong *et al* 2012) and reduce apoptosis following cold injury (Yi *et al* 2007). However, cold hardening also carries costs in terms of future reproductive output (Overgaard *et al* 2007; Everman *et al* 2018), suggesting that avoiding cold hardening could be beneficial if temperatures never drop to lethally cold. Individual *D. melanogaster* genotypes vary in their cold hardening response (Gerken *et al* 2015, 2018); therefore, different cold hardening abilities may be advantageous under different conditions. In addition to genetic influences, specific aspects of environmental exposure, such as the temperature and duration of cold exposure (Cjazka and Lee 1990; Kelty and Lee 1999) also affect cold hardening. We thus hypothesized that the cold hardening response might vary seasonally in temperate climates via some combination of genetic adaptation and environmental influences.

In the present study, we ask whether the cold hardening response in *D. melanogaster* varies seasonally in field-reared populations, and if so, whether this variation occurs as a result of environmental influences, genetic changes, or some combination thereof. Although other studies have examined the occurrence of rapid cold hardening in the field (e.g. Kelty 2007; Overgaard and Sorensen 2008), in this study we measure the effect of seasonal changes on the response to a consistent cold hardening treatment. Over the course of multiple seasons in a single year, we collected flies from outdoor mesocosms and then subjected them to a controlled pre-exposure to cool temperatures followed by a freeze tolerance test. By contrasting outdoor-caught flies, their lab-reared offspring, and their grandchildren to similarly treated flies that were reared entirely indoors, we found that cold hardening does not evolve over seasonal timescales but shows dramatic season-specific plasticity. We further show that this plasticity is potentially caused by thermal exposure in the field and is likely influenced by nutrition, age, and other factors. Taken together, our work suggests that cold hardening is a highly plastic trait that does not exhibit classic signatures of adaptive tracking.

## Methods

### The hybrid swarm

Using inbred fly lines from various regions of the Eastern United States and the Bahamas, we created two outbred and genetically diverse populations of flies (populations A and B) for use in our experiments. We created each population by crossing a total of 34 fly lines representing each of our chosen regions, with the particular lines from each region picked at random. We used flies from Bowdoinham, Maine (NCBI BioProject # PRJNA383555); Ithaca, New York (Grenier *et al* 2015), spring and fall collections from Linvilla Orchard, Media, Pennsylvania (Behrman *et al* 2018), Raleigh, North Carolina (MacKay *et al* 2012), the Southeastern United States (Kao *et al* 2015), and the Bahamas (Kao *et al* 2015). The initial crosses were established with four sets of 34 round-robin crosses. After each population was established, we maintained them with two-week, non-overlapping generations in indoor mesh cages (2m x 2m x 2m; Bioquip product number 1406C) at a population size of approximately 10,000 flies per generation. Each generation received approximately 5 L of standard cornmeal-molasses media sprinkled with live baker’s yeast. We reared the flies in the lab for approximately 32 generations before transferring subsets of each population to outdoor cages in June 2018.

### Pretreatment

We pretreated flies by placing them in a temperature-controlled chamber at approximately 11 °C with a 9L:15D photoperiod for 13-15 days (typically 14 days; longer or shorter precooling periods occasionally occurred). We hereafter refer to this pretreatment period as precooling. During precooling, we held flies in vials containing cornmeal-molasses food. Vials contained 25 male flies each except in rare circumstances when fewer flies were available. We sorted flies using CO_2_ sedation. We sedated flies for a maximum of twenty minutes, which is well under the sedation duration found to impact the cold hardening response (Nilson *et al* 2006).

### The freeze assay

Due to the high level of thermal sensitivity involved in *D. melanogaster*’s cold tolerance (Czajka and Lee 1990), we designed the freeze assay with the goal of creating the most consistent thermal environment possible both during and between freeze assays. We logged air and water temperature with EL-WiFi-DTP+ (dataq.com) temperature probes and the manufacturer’s EasyLog software. By placing the Falcon tubes that contained the flies inside a saltwater (~3 M NaCl) bath inside a freezer, rather than in the freezer directly, we reduced temperature fluctuations experienced by the flies. Air temperature in the freezer fluctuated by several degrees Celsius, whereas temperature in the saltwater bath varied on the scale of tenths of a degree (**Figure 1)**.

**Figure 1.**
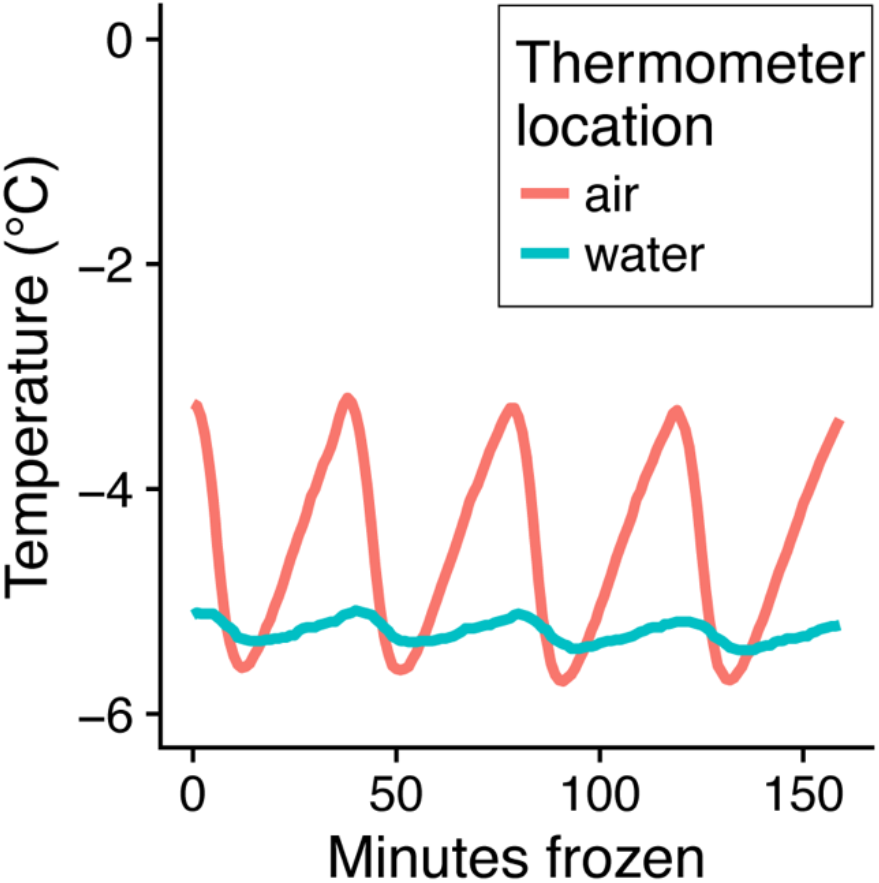
A saltwater bath provides a stable thermal environment during the freeze assay. Temperature data recorded during a representative freeze assay. Temperature probes exposed to air inside the freezer recorded temperature fluctuations ranging from approximately −3.3 °C to −5.7 °C over the course of the freeze assay. Temperature probes submerged in a saltwater bath inside the freezer recorded temperature fluctuations ranging from approximately −5.1 °C to −5.4°C.

Following precooling, we noted the number of flies that had died during the precooling process prior to conducting the freeze assay. We transferred each vial of flies to a 5 mL snap-cap Falcon tube and suspended the tubes in the salt solution held at approximately −5 °C within a chest freezer. We used weighted blocks to keep the tubes submerged in liquid up to the rim of the cap. In order to minimize temperature fluctuations in the water bath, we added tubes into the bath in small groups for each time point and removed all the tubes at the end of the assay. At the conclusion of the freeze, we transferred flies into their original vials containing food and held the food vials upside down so that unconscious flies would not become stuck in the food. The next day, we recorded the number of survivors or the number dead within each vial (whichever number was smaller). Flies that exhibited the ability to stand stably and walk were considered to be alive, while flies that were immobile, or flies that exhibited spastic motions such as twitching but were not stable in their movements and stance, were considered dead following similar definitions of Czajka and Lee (1990) and Gerken *et al* (2015).

### Seasonal assay

Field studies were conducted at Morven Farms in Charlottesville, VA (37°58’02.9”N, 78°28’26.4”W). We established outdoor cages (2m x 2m x 2m) surrounding peach trees and initiated each population with approximately 3,000 flies from our indoor hybrid swarm populations. Cages 1-6 were started on June 2^nd^, 2018, and cages 7-11 were started on September 11^th^, 2018. Cages 1-6 were fed with bananas and apples (approximately 3.6 kg of each added to the cages weekly until November, when feeding was done biweekly) and were initially inoculated with live yeast. Cages 7-11 were fed weekly with 2.5 L of cornmeal-molasses fly food, the same food that fed the indoor cages. We collected “F0” flies from the outdoor cages using nets or aspirators. Note that although these flies are actually advanced generations of the outbred populations, we refer to them as F0s for the given collection point. On the same day, we collected F0 flies from the indoor cages. We subjected males from the indoor and outdoor cages to the precooling and freeze assays as described above. We used only males because female *D. melanogaster* cannot be readily distinguished from another common inhabitant of orchards, *D. simulans,* based on external morphology (Markow and O’Grady 2006). Our outdoor cages were not completely impenetrable to other insect species, and *D. simulans* males were occasionally found in our collections.

In the lab, we established 25 isofemale lines from each cage in vials. We screened the offspring of the isofemale lines for presence of *D. simulans* and discarded these lines (on average, fewer than 10/150 lines per collection time point). We combined all *D. melanogaster* F1s together to collect males for the freeze assay and to establish F2s. F1 males collected from the isofemale lines were precooled, frozen, and assayed for survival. The F1 flies were propagated in bottles and the F2 males were precooled, frozen, and assayed for survival. Heavy rain limited the 7/24/18 collection; we obtained sufficient F0 females to establish lines but not enough males to generate a freeze survival curve. All F1 isofemale lines from the 11/7/18 and 11/21/18 collections were lost in an incubator failure, so we were unable to assay the offspring of these collections. We collected again on 11/30/18 to replace these samples but did not perform a freeze assay on the 11/30/18 F0s.

We downloaded weather data from a station located at Carter Mountain, which is approximately 2 km from our field site, to obtain information on average daily temperatures during our experiments (https://www.wunderground.com/dashboard/pws/KVACHARL73).

### Nutrition and antimicrobial assays

We made standard cornmeal-molasses fly food in a small batch using a hot plate and added standard amounts of Tegosept and propionic acid to half the batch (**Table 1**). We used Agricor Fine Yellow Cornmeal and Golden Barrel Blackstrap Molasses in our cornmeal-molasses fly food. In a separate batch of food, we substituted the cornmeal and molasses by volume with pureed bananas (**Table 1**). We added standard concentrations of Tegosept and propionic acid to half of this batch as well. We allowed flies from each hybrid swarm cage to lay eggs in bottles of the different food types. After these eggs matured into adult flies, we collected males from all treatments and placed them in vials containing standard cornmeal-molasses fly food, precooled them, and subjected them to the freeze assay. Therefore, the developmental nutritional environment varied, but adult nutrition during the precooling period was identical across all assays.

**Table 1:**
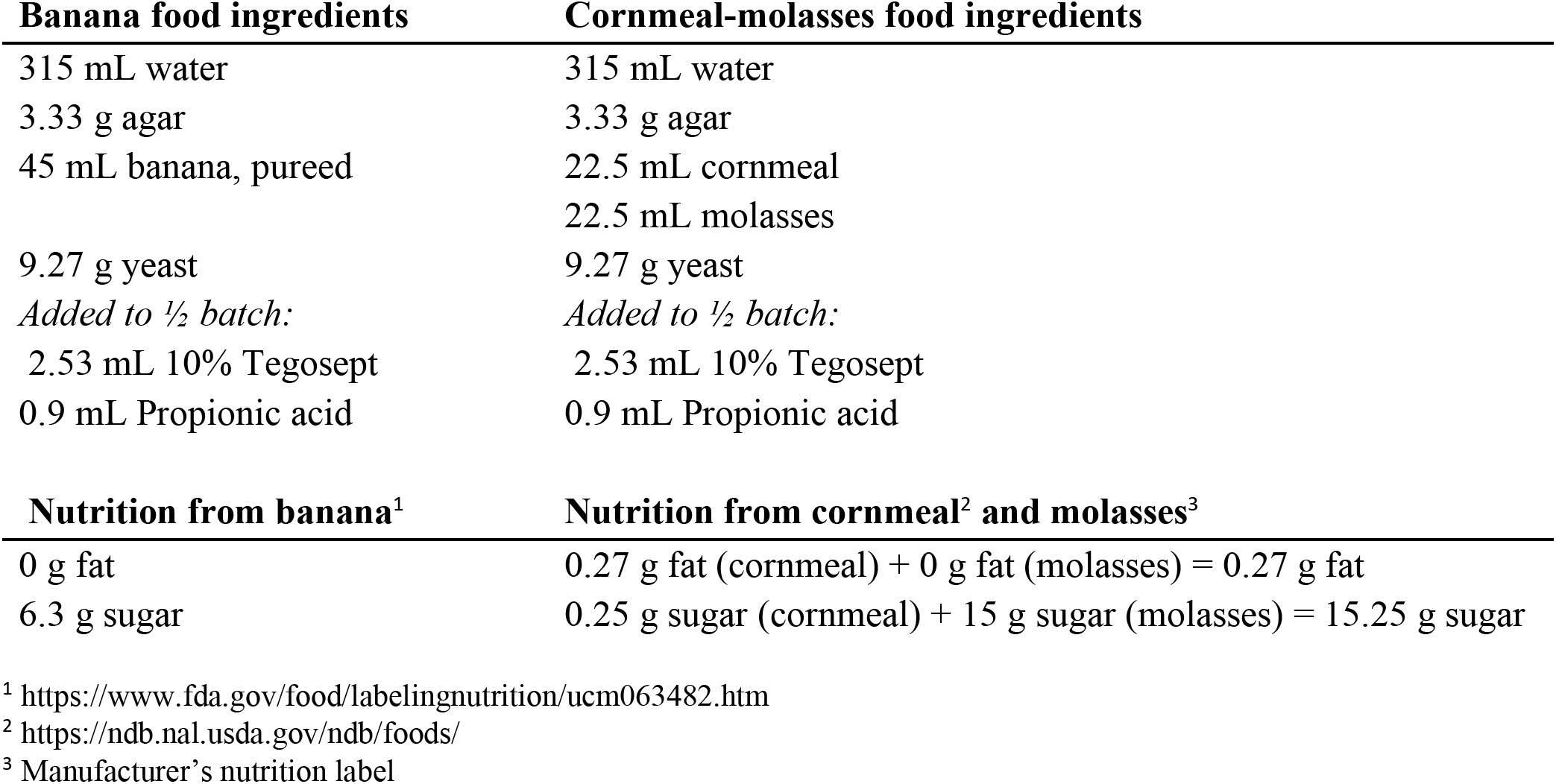
Ingredient list and nutritional analysis of laboratory fly food

### Supplemental yeast assay

We collected eggs from indoor hybrid swarm cages in bottles with cornmeal-molasses fly food. We added live bakers’ yeast to the surface of half of the bottles and continued to supplement these bottles with yeast throughout the development of the flies. We collected adult males from each treatment and placed them in vials with standard fly food and no live yeast. We precooled them for two weeks and subjected them to the freeze assay.

### Age assay

We collected embryos from the indoor hybrid swarm cages at two-week intervals for six weeks and reared them to adulthood. We passaged the adult flies to fresh food weekly to prevent eclosion of new adults. We collected adult males once the youngest of the cohorts had eclosed, and we then precooled, froze, and measured survival of flies from all three age cohorts in a single assay.

### Density assay

We based our density assay on a previous study (Henry *et al* 2018). We used four density levels: approximately 5 embryos/mL, 40 embryos/mL, 120 embryos/mL, and 300 embryos/mL of fly food. We collected embryos from the indoor cages on cornmeal-molasses fly food plates and counted embryos into vials containing 2 mL of cornmeal-molasses fly food. As higher density vials took longer to develop, we waited to collect adults until every vial had sufficiently eclosed. As a result, the flies in higher density vials were several days younger than the flies in lower density vials at the time of precooling. We collected adult males and subjected them to the precooling and freeze assay.

### Analysis

We analyzed our results using R (version 3.4.2; R Core Team 2017). We used packages *data.table* (Dowle and Srinivasan 2017), *foreach* (Microsoft and Weston 2018), *ggplot2* (Wickham 2009), and *cowplot* (Wilke 2017) for data manipulation, looping, and graphing. We used *lubridate* (Grolemund and Wickham 2011) for date conversions. We calculated LT50 using the *dose.p()* function from the *MASS* package (Venables and Ripley 2002). All general linear model results are summarized with an ANOVA. We used *lme4* (Bates *et al* 2015) to create a generalized linear mixed-effects model (glmm) where noted.

## Results

### Effects of seasonal exposure on cold hardening

To test the hypothesis that seasonal conditions influence cold hardening, we collected monthly samples of outbred flies reared in fruit-fed outdoor mesocosms and cornmeal-molasses-fed laboratory cages and assessed their cold tolerance following two weeks of precooling in the lab. We used the resulting survival curves to calculate the time required to kill 50% of the flies (hereafter, “LT50”). From June until early November, the LT50 of fruit-fed, outdoor F0 flies was significantly lower than the LT50 of indoor F0 flies (**Figure 2A; Table 2**). However, in late November, the outdoor F0 flies had an LT50 greater than that of the indoor F0 flies (**Figure 2A; Table 2)**. We collected outdoor flies on December 10^th^, 2018, but the sample size was insufficient to generate a survival curve. However, the limited data matched the trend from the November collection; after being frozen for 164 minutes, the survival for the outdoor flies was 84% while the survival for the indoor flies was 72% (Fisher’s exact test, *P* = 0.39).

**Figure 2.**
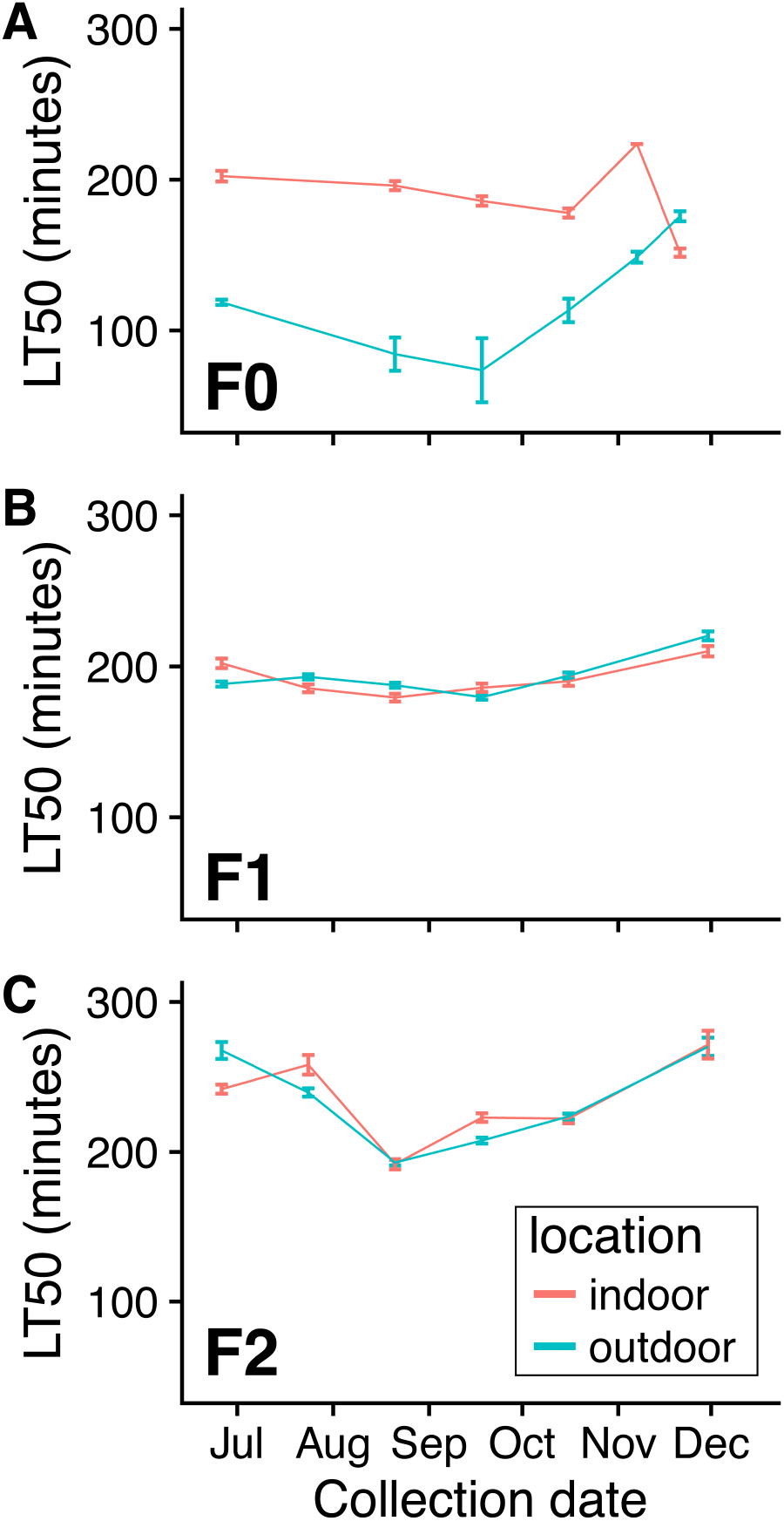
Seasonal plasticity, but not adaptive tracking, in the cold hardening response. Cold-hardened freeze tolerance for flies collected from indoor and outdoor cages and their offspring. We calculated the LT50, or the time during the freeze assay at which 50% of flies perished, using a general linear model. **A)** The F0 generation was collected directly from the indoor or outdoor cages. In the F0 generation, we observed significantly lower LT50s for outdoor flies relative to indoor flies during the summer and fall seasons (**Table 2**). However, in the late November collection, we observed an increased LT50 for outdoor flies compared to indoor flies (*P* = 5.31 x 10^−7^). **B&C)** The F1 and F2 generations were reared in laboratory conditions. We generally did not observe differences between LT50 values for outdoor and indoor flies in the F1 and F2 generations, regardless of collection time (**Table 2**). Error bars represent standard error of the LT50. Standard error for F0 indoor data from November 7^th^, 2018 is set to zero for clarity (SE = 2927.8 due to incomplete survival curve).

**Table 2.** General linear mixed-effects model comparisons of freeze tolerance of outdoor and indoor flies across collection points and generations.

| Generation | Collection date | *P*-value | Direction |
| --- | --- | --- | --- |
| F0 | 6/26/2018 | **2.17 x 10<sup>-31</sup>** | indoor greater |
| F0 | 8/21/2018 | **4.41 x 10<sup>-23</sup>** | indoor greater |
| F0 | 9/18/2018 | **1.40 x 10<sup>-16</sup>** | indoor greater |
| F0 | 10/16/2018 | **3.44 x 10<sup>-14</sup>** | indoor greater |
| F0 | 11/7/2018 | **7.42 x 10<sup>-14</sup>** | indoor greater |
| F0 | 11/21/2018 | **5.31 x 10<sup>-7</sup>** | outdoor greater |
| F1 | 6/26/2018 | **1.33 x 10<sup>-4</sup>** | indoor greater |
| F1 | 7/24/2018 | 0.162 | - |
| F1 | 8/21/2018 | 0.056 | - |
| F1 | 9/18/2018 | 0.106 | - |
| F1 | 10/16/2018 | 0.562 | - |
| F1 | 11/30/2018 | 0.082 | - |
| F2 | 6/26/2018 | 0.022 | - |
| F2 | 7/24/2018 | 0.027 | - |
| F2 | 8/21/2018 | 0.853 | - |
| F2 | 9/18/2018 | **0.003** | indoor greater |
| F2 | 10/16/2018 | 0.833 | - |
| F2 | 11/30/2018 | 0.099 | - |

Bold text indicates tests that pass Bonferroni correction. Direction indicates which population had greater LT50 (higher freeze tolerance).

In the field experiment, we observed that the effects of the environment prior to precooling persisted through the two-week precooling period. In order to test whether the thermal environment experienced by the flies before precooling influenced their cold hardening, we tested for a relationship between the average temperature on the day of collection and the difference in the cold hardening response of outdoor and indoor F0 flies. We observed a significant negative correlation between the average temperature on the day of collection and the difference in LT50 (**Figure 3**; linear model; R^2^ = 0.90, *P* = 0.0024). Therefore, as temperatures became colder, the cold hardened freeze tolerance increased linearly for the outdoor F0 flies relative to the indoor flies. The regression is also significant when the coldest collection is excluded (R^2^ = 0.81, *P* = 0.02), suggesting that prior exposure affects the cold hardening response even at moderate to warm temperatures.

**Figure 3.**
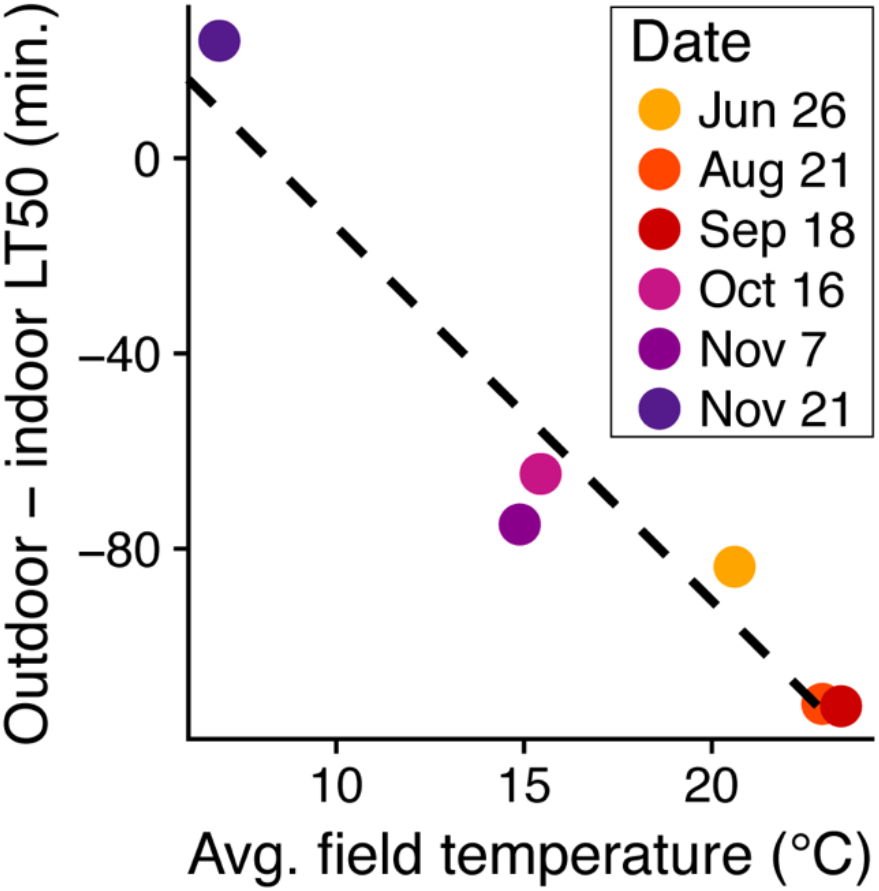
Improved cold hardening with decreasing field temperatures. Temperature on day of collection is negatively correlated with the difference in LT50 values for outdoor and indoor F0 flies (linear model; R^2^ = 0.90, *P* = 0.002).

To test for transgenerational effects of seasonal exposure on the cold hardening response, we compared the lab-reared F1 offspring of flies collected indoors to the lab-reared F1 offspring of flies collected outdoors in a common garden assay. We observed that the cold hardening response was generally consistent between the lab-reared offspring of flies collected from indoor and outdoor cages (**Figure 2B, Table 2**). The similarity between indoor and outdoor F1 flies suggests that the differences in the cold hardening responses in the F0 flies were not passed on to their offspring. We note that although indoor and outdoor LT50s were significantly different for the 6/26/18 collection (**Table 2**), we did not observe a consistent difference or pattern in indoor versus outdoor F1 cold hardening responses.

We tested for genetic changes in the cold hardening response by examining the lab-reared F2 offspring of flies from indoor and outdoor cages. As in the F1s, we also observed little difference in the cold hardening response between outdoor and indoor flies (**Figure 2C; Table 2**). We note that the difference in LT50 was significant for the 9/18/18 collection, but again we did not observe a consistent difference or pattern between the indoor and outdoor F2 cold hardening responses. Therefore, our data do not provide evidence of seasonal genetic changes in cold hardening. Although we observed some differences in the overall cold hardened freeze tolerance of F2 flies across collection points (**Figure 2C;** compare July and August), these seasonal changes occurred in parallel between the indoor and outdoor populations, so we attribute this pattern to experimental artifact and not seasonal evolution. This result emphasizes the invaluable nature of internal controls—in this case, the indoor cages—when conducting seasonal experiments.

Surprisingly, we observed that F2 flies were more freeze tolerant following cold hardening than their F1 parents for some collections (**Figure 2B-C**, compare July F1s and F2s). Differences in rearing conditions between the generations may have contributed to the differences in cold hardening: F1 flies were reared in vials, whereas F2 flies were reared in bottles. We investigated whether container type was a potential cause by rearing the 9/18/18 set of F1 flies in both bottles and vials. We did not observe significant differences in the cold hardening response resulting from differences in container type (**Table 3**; *P* = 0.191), though vial-reared flies had a slight increase in the cold hardening response compared to bottle-reared flies. We suggest that differences in freeze tolerance between the F1 and F2 generations may have been caused by stochastic differences in rearing conditions in the F2 bottles as compared to the relatively consistent conditions of F1 isofemale lines.

**Table 3.** General linear models of the effects of experimentally manipulated rearing conditions on the cold hardening response.

| **Figure** | **Factor** | **Df** | **Sum sq** | **Mean sq** | **F** | **P** |
| --- | --- | --- | --- | --- | --- | --- |
| **N/A** | Minutes frozen | 1 | 315.19 | 315.19 | 519.36 | **< 2 x 10<sup>-16</sup>** |
|  | Bottle vs vial | 1 | 1.06 | 1.06 | 1.75 | 0.19 |
|  | Indoor vs outdoor | 1 | 0.20 | 0.20 | 0.34 | 0.56 |
|  | Cage | 6 | 1.12 | 0.19 | 0.31 | 0.93 |
|  | Residuals | 68 | 41.27 | 0.61 |  |  |
| **4A** | Minutes frozen | 1 | 20.82 | 20.82 | 26.04 | **1.61 x 10<sup>-4</sup>** |
|  | Indoor vs outdoor | 1 | 21.17 | 21.17 | 26.48 | **1.49 x 10<sup>-4</sup>** |
|  | Cage | 6 | 1.63 | 0.27 | 0.34 | 0.91 |
|  | Residuals | 14 | 11.20 | 0.80 |  |  |
| **4B** | Minutes frozen | 1 | 114.36 | 114.36 | 328.59 | **< 2 x 10<sup>-16</sup>** |
|  | Indoor vs outdoor | 1 | 0.70 | 0.70 | 2.00 | 0.17 |
|  | Cage | 5 | 2.19 | 0.44 | 1.26 | 0.30 |
|  | Residuals | 34 | 11.83 | 0.35 |  |  |
| **5A** | Minutes frozen | 1 | 118.40 | 118.40 | 132.06 | **1.08 x 10<sup>-14</sup>** |
|  | Food substrate | 1 | 51.74 | 51.74 | 57.71 | **1.78 x 10<sup>-9</sup>** |
|  | Antimicrobials | 1 | 0.00 | 0.00 | 0.005 | 0.94 |
|  | Cage | 1 | 0.14 | 0.14 | 0.16 | 0.69 |
|  | Residuals | 43 | 38.55 | 0.90 |  |  |
| **5B** | Minutes frozen | 1 | 18.04 | 18.04 | 30.58 | **5.54 x 10<sup>-4</sup>** |
|  | Supplemental yeast | 1 | 5.31 | 5.31 | 9.01 | **0.017** |
|  | Cage | 1 | 3.18 | 3.18 | 5.39 | **0.049** |
|  | Residuals | 8 | 4.72 | 0.59 |  |  |
| **5C** | Minutes frozen | 1 | 31.73 | 31.73 | 26.20 | **1.82 x 10<sup>-5</sup>** |
|  | Age | 2 | 86.45 | 43.22 | 35.70 | **1.52 x 10<sup>-8</sup>** |
|  | Cage | 1 | 0.44 | 0.44 | 0.367 | 0.55 |
|  | Residuals | 29 | 35.12 | 1.21 |  |  |
| **5D** | Minutes frozen | 1 | 40.49 | 40.49 | 50.37 | **8.09 x 10<sup>-6</sup>** |
|  | Density | 2 | 12.34 | 6.17 | 7.68 | **0.0063** |
|  | Cage | 1 | 0.00 | 0.00 | 0.00 | 0.99 |
|  | Residuals | 13 | 10.45 | 0.80 |  |  |

Note: tests conducted only on timepoints with complete data.

### Plastic effects of nutrition on cold hardening

In addition to experiencing different thermal environments, the indoor and outdoor flies described above consumed different foods prior to cold hardening, which led to the hypothesis that differences in nutritional intake might also affect cold hardening in the seasonal experiments. In October, we collected F0 flies from both the cornmeal-molasses-fed outdoor cages and the fruit-fed outdoor cages within one day of each other. While the fruit-fed outdoor flies exhibited a cold hardening response significantly lower than that of the indoor flies (**Figure 4A; Table 3;** *P* = 1.49 x 10^−4^), the cornmeal-molasses-fed outdoor flies exhibited comparable freeze tolerance to the indoor flies (**Figure 4B**; **Table 3**; *P* = 0.17). Therefore, flies that experienced nearly identical outdoor thermal regimes greatly differed in the cold hardening response depending on their nutritional exposure. Notably, the majority of flies in the cornmeal-molasses-fed outdoor cages died prior to our next collection at the end of November, despite their apparent enhanced cold hardening ability. The mass mortality of these cages may have been due to the absence of thermal refugia (rotting fruit) during subfreezing temperatures in late fall.

**Figure 4.**
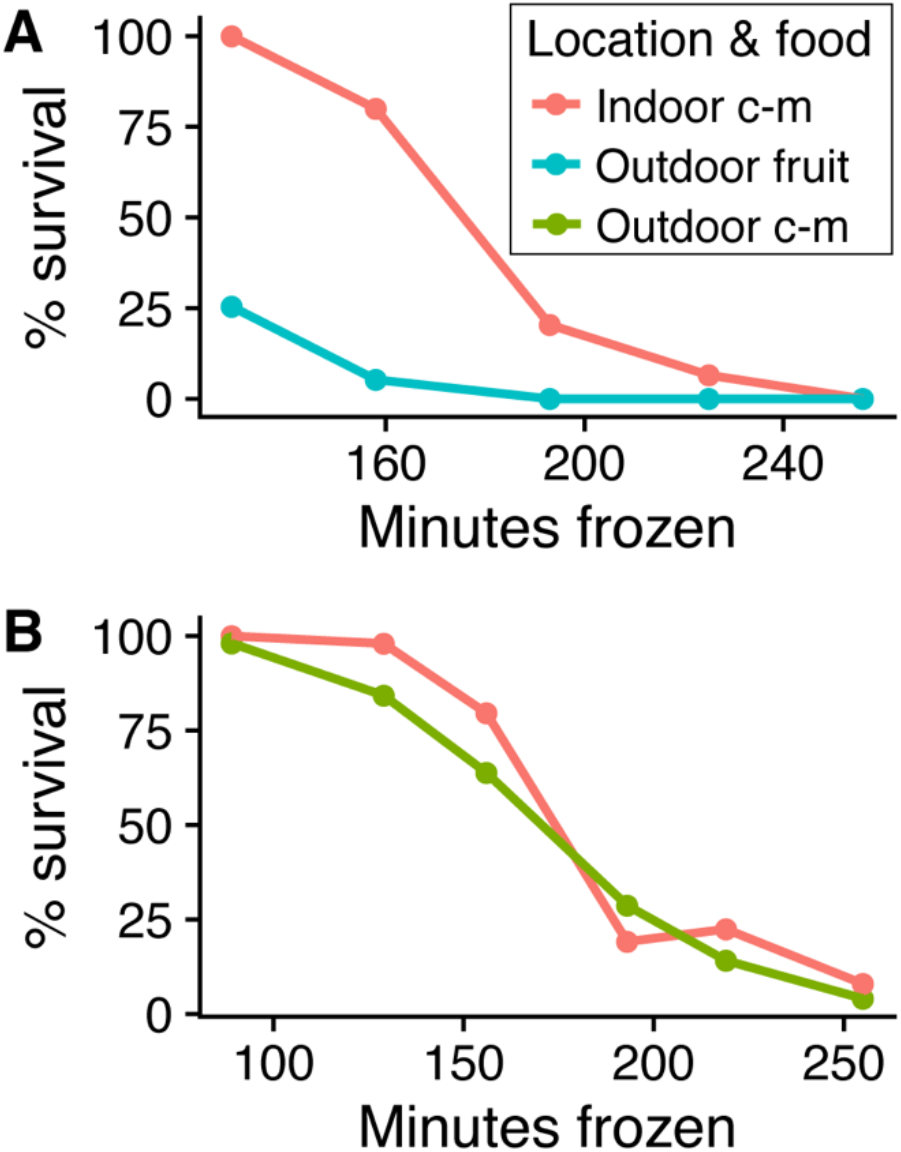
The cold hardening response varies between outdoor flies fed different substrates. Cold hardened freeze survival curves for F0 flies collected from fruit-fed cages on October 16^th^, 2018 (**A**) and cornmeal-molasses (c-m) fed cages October 15^th^, 2018 (**B**) relative to indoor controls collected on each day. Although outdoor flies from both food treatments experienced comparable thermal conditions, we observed a significant difference in cold hardened freeze tolerance for indoor flies versus outdoor fruit-fed flies (*P* = 1.49 x 10^−4^, **A**; **Table 3**) but not for indoor flies versus outdoor cornmeal-molasses-fed flies (*P* = 0.17, **B**; **Table 3**). Outdoor survival curves are pooled from six cages in A and five cages in B. Indoor survival curves are pooled from two cages.

We tested several variables that could explain the differences in cold hardening in F0 flies reared on different foods under similar thermal conditions. First, we compared the cold hardening response of indoor flies reared on either a fruit substrate (banana-based) or the cornmeal-molasses substrate. We observed that flies reared on banana-based food exhibited a decreased cold hardening response relative to flies reared on cornmeal-molasses food (**Figure 5A; Table 3**; *P* = 1.78 x 10^−9^). We also tested whether adding antimicrobials influenced cold hardening, since the cornmeal-molasses food contained Tegosept and propionic acid while the rotting fruit did not. We did not observe an effect of antimicrobial presence on cold hardening (**Figure 5A, Table 3**; *P* = 0.95). Therefore, we suggest the cornmeal-molasses diet improved the indoor F0 cold hardening response relative to outdoor flies during summer months (**Figure 2A**) and also improved the cold hardening response of outdoor flies fed cornmeal-molasses food (**Figure 4**).

**Figure 5.**
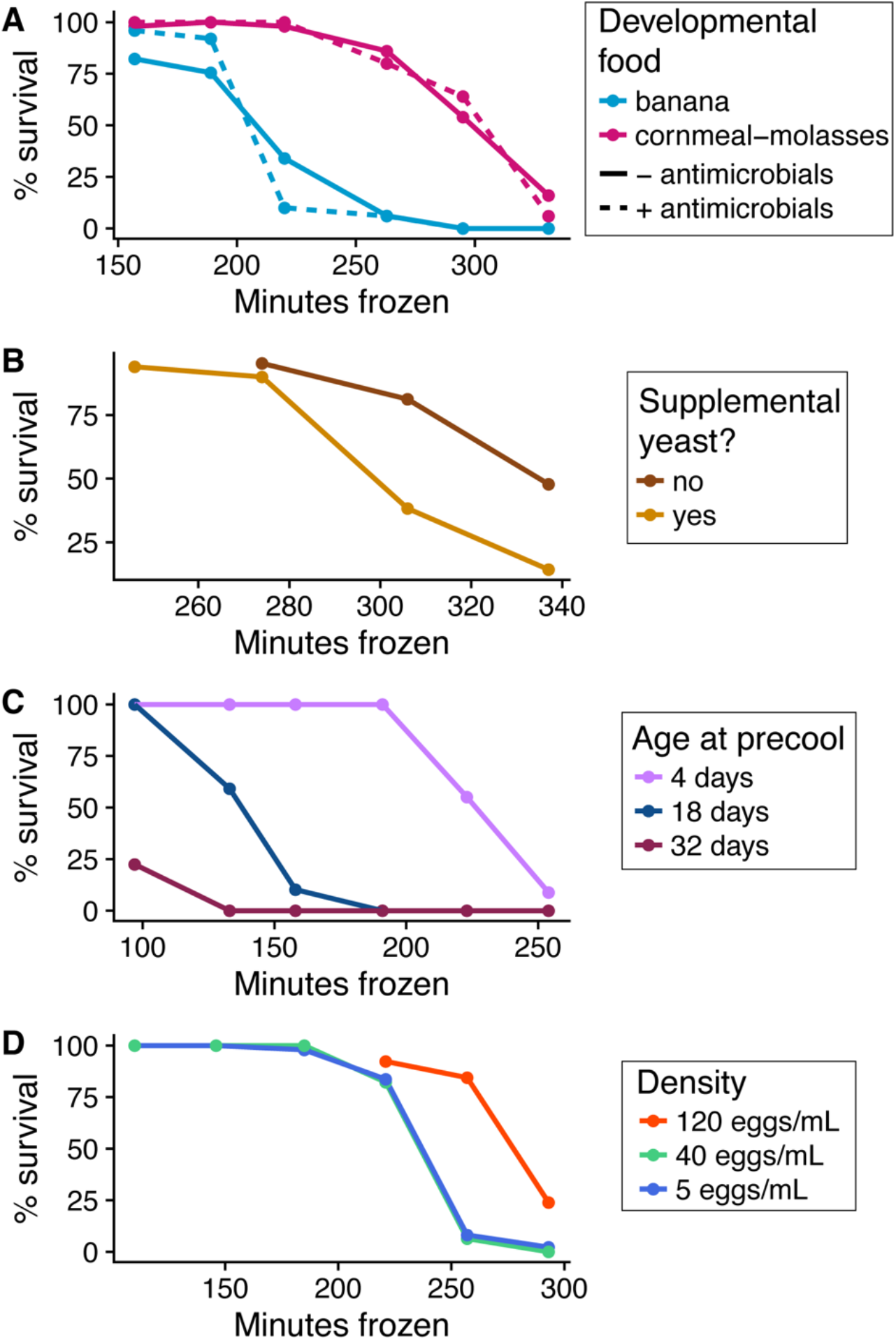
Life history and nutrition influence the cold hardening response. **A)** Cold hardened freeze survival curves for indoor flies reared on banana-based or cornmeal-molasses food, with or without antimicrobials added. The cold hardening response is higher for flies reared on cornmeal-molasses food compared to flies reared on banana-based food (**Table 3**; *P* = 1.78 x 10^−9^). No effect of antimicrobials was observed (P = 0.95). **B)** Cold hardened freeze survival curves for flies reared with or without supplemental yeast. Flies supplemented with yeast exhibited a slightly decreased cold hardening response (**Table 3**; *P* = 0.017). **C)** Cold hardened freeze survival curves of indoor flies of varying ages. Older flies exhibited a lower cold hardening response, while younger flies exhibited a higher cold hardening response (**Table 3**; *P* = 1.52 x 10^−8^). **D)** Cold hardened freeze survival of flies reared under varying larval density conditions. Flies developed in low and mid density conditions (5 eggs/mL, 40 eggs/mL) had similar cold hardening responses. Flies developed in high density conditions (120 eggs/mL) exhibited a greater cold hardening response than low and mid density flies (**Table 3**; *P* = 0.0063). All survival curves are pooled from two replicates: one from cage A and one from cage B.

We also hypothesized that yeast availability could contribute to differences in cold hardening ability. Specifically, the cornmeal-molasses fly food contained yeast as an ingredient, while the cages with apples and bananas relied on yeast growth following an inoculation plus any naturally occurring yeasts. We observed a slight decrease in cold hardening ability for flies supplemented with extra yeast (**Figure 5B**; **Table 3**; *P* = 0.017). Though we cannot directly quantify yeast availability in the outdoor cages, seasonal variation in yeast growth may have had a minor influence on cold hardening in the flies in the fruit-fed outdoor cages.

### Plastic effects of life history on cold hardening

The indoor flies were maintained on a two-week generation cycle, while the outdoor flies were able to breed in overlapping generations, likely resulting in a more complex age structure in the outdoor cages. To determine whether age differences could explain the indoor-outdoor differences, we tested the cold hardening response of lab-reared flies of various ages. We observed that younger flies had greater cold hardening responses than older flies (**Figure 5C; Table 3;** *P* = 1.52 x 10^−8^). Although differences in age structure could potentially explain the lower cold hardening exhibited by fruit-fed outdoor F0 flies as compared to indoor F0 flies, we expect that age structure between the fruit-fed and cornmeal-molasses-fed outdoor cages (shown in **Figure 4**) should be similar. If age structure alone was causing the outdoor F0 flies to have a lower cold hardening response in the summer and fall, we would expect the cornmeal-molasses-fed outdoor cages to also have decreased cold hardening, which was not the case.

Density is a final possibly causal factor in the indoor-outdoor cold hardening differences, since different food substrates and different age structures could lead to different larval densities. We reared larvae at varying densities and observed an increase in cold hardening ability at relatively high densities only (**Figure 5D; Table 3;** *P* = 0.0063). We suggest that differences in density may have contributed to the observed cold hardening differences in the outdoor and indoor cages; however, factors such as thermal environment and nutrition are more compelling causal factors given our experimental results.

## Discussion

Many phenotypes undergo seasonal fluctuations in response to the varying demands of temperate environments. Some organisms exhibit phenotypic plasticity, meaning that an environmental stimulus induces a change in the phenotype. Some populations adaptively track, meaning that some genotypes are more favorable in a given season, and therefore, individuals having those genotypes will be more abundant during that season. In our study, we asked whether the cold hardening response in *D. melanogaster* varies seasonally and whether such variation is a product of plasticity or adaptive tracking. We found that the cold hardening response increases as outdoor temperature decreases at the onset of winter. We also determined that, while cold hardening is highly plastic, the trait does not undergo seasonal evolution. Therefore, we conclude that seasonal fluctuations in the cold hardening response are governed by environmental and developmental variables rather than adaptive tracking.

### The cold hardening response varies seasonally

Previous studies have demonstrated that cold hardening occurs under natural conditions in *D. melanogaster* using field studies (Kelty 2007; Overgaard and Sorensen 2008). These studies placed flies outdoors and then measured their cold tolerance following the natural cold hardening treatment. In our work, we further exposed flies to a consistent precooling treatment after they were exposed to field conditions. This experimental design allowed us to elucidate the effects of field exposure that persisted through a consistent, controlled cold hardening regime. We found that the onset of winter conditions correlated with an increased cold hardening response for outdoor flies (**Figures 2A, 3**). Previous data have shown that the cold tolerance of flies kept outdoors for several hours or days correlates negatively with outdoor temperatures (Overgaard and Sorensen 2008). We have demonstrated that the effect of field conditioning either persists through two weeks of precooling or modulates the ability to cold harden in laboratory conditions.

We cannot be certain of the exact influence of field exposure on our results because limited population sizes prevented us from testing the basal cold tolerance of each seasonal collection in addition to the cold hardened cold tolerance. On one hand, winter conditions prior to the laboratory precooling treatment could simply serve to extend the precooling period, producing a stronger cold hardening response. On the other hand, winter conditions may induce a plastic change in the *ability* to cold harden, thereby enhancing the cold hardening that occurred during the laboratory precooling treatment. We suggest that the former is the more plausible explanation given knowledge from previous studies. Longer precooling periods result in a greater increase in cold tolerance (Cjazka and Lee 1990) and repeated exposure to cold has an additive effect on cold tolerance (Kelty and Lee 2001). Therefore, we suggest that flies sampled during cold periods experienced extra cold hardening prior to being brought into the laboratory, and thus their cold tolerance was enhanced.

### The cold hardening response does not evolve over seasons

Two possible reasons could explain why we did not observe adaptive tracking of the cold hardening response (**Figure 2B&C; Table 2**). One possible explanation is that our outbred populations of *D. melanogaster* carried limited heritable variation in cold hardening. However, two studies have demonstrated heritable variation in this trait in flies from North Carolina, a population that was included in our hybrid swarms (Gerken *et al* 2015, 2018). Thus, we suggest that the absence of adaptive tracking in cold hardening is not due to a lack of genetic variation for this trait within the experimental population.

A second possibility is that the cold hardening response is not subject to local adaptation over space and time. Several lines of evidence from previous work are consistent with this model. Field studies have tested for evidence of natural selection in cold hardening in wild populations of *D. melanogaster.* Flies native to tropical or temperate regions exhibit a similar capacity to cold harden despite differences in basal cold tolerance, suggesting that cold hardening undergoes minimal evolution across latitudinal clines (Hoffmann and Watson 1993). Similarly, flies native to temperate or tropical climates also display comparable overwintering survival in a common environment (Mitrovski and Hoffmann 2001), suggesting that their cold acclimation ability may be similar. Finally, wild-caught temperate flies and their recent descendants have been shown to exhibit comparable cold hardening ability to laboratory flies, implying that experiencing natural conditions did not select for changes in cold hardening (Kelty 2007). Given these results, the lack of evolution of the cold hardening response in the outdoor cages is not surprising.

Although we did not observe adaptive tracking in cold hardening, this absence of seasonal evolution is not shared by all cold-related traits. For example, the timing of winter reproduction varies between flies originating from tropical and temperate regions (Mitrovski and Hoffmann 2001; Hoffmann *et al* 2003), and seasonal and latitudinal variation occurs in diapause propensity (Schmidt *et al* 2005, Schmidt and Conde 2006). A study of diapause induction in the same outdoor cages studied here demonstrated evolution of increased propensity for diapause induction in late fall (Erickson *et al*, in prep), suggesting that this population did in fact carry heritable variation for overwintering-related traits. Notably, we observed obvious decreases in the population sizes of our outdoor cages in late fall, suggesting that selection was indeed happening. However, since this population reduction did not affect the average cold hardening response in subsequent generations, we suggest that the selection on the population did not involve cold hardening.

### Cold hardening is plastic and modified by a variety of conditions

The cold hardening response varies seasonally; however, we did not observe evidence of adaptive tracking. Combined with correlations between the cold hardening response and outdoor temperature (**Figure 3**), we conclude that the seasonal variation observed was a result of plasticity. However, the thermal environment is likely not the sole stimulus that triggers plastic changes in the cold hardening response.

#### Nutrition partially explains differences in cold hardening between indoor and outdoor flies

We demonstrated that the cold hardening response of flies reared on banana-based food is significantly less than the cold hardening response of flies reared on cornmeal-molasses food (**Figure 5A; Table 3**). The cornmeal-molasses food contained slightly more fat and more than double the sugar of the banana food (**Table 1**). A number of studies have examined the effects of nutritional profiles, particularly fats and sugars, on cold hardening ability.

The role of fats in cold hardening could be mediated by a number of physiological processes. Cold hardening typically changes the makeup of lipid membranes (Overgaard *et al* 2005; Overgaard *et al* 2006; but see MacMillan *et al* 2009). Additionally, increasing cholesterol consumption during larval development has been shown to enhance both baseline cold tolerance and the cold hardening response (Shreve *et al* 2007). Although there is little difference in fat content between our foods, increasing dietary sugars in flies can result in greater fat storage (Colinet *et al* 2013). Furthermore, differences in the specific fats found in each food could affect cold hardening; the availability of dietary fats can impact the process of desaturation or other alterations to lipid composition (Overgaard *et al* 2005).

Dietary sugar itself may also have a direct effect on the cold hardening response. Certain sugars, particularly glucose and trehalose, are increased in the fly following cold hardening, and greater increases in sugars correspond with a greater cold hardening response (Overgaard *et al* 2007; but see evidence of dietary sugar lowering basal cold tolerance in Colinet *et al* 2013). Flies fed increased levels of sugar, and so possessing greater sugar stores, may be able to elicit a greater cold hardening response either directly or via changes in fat storage. Taken together, sugar serves as a potential mediator of the increased cold hardening response in flies fed cornmeal-molasses food as compared to banana food.

Live yeast is another nutritional factor that may have contributed to plastic differences in cold hardening between indoor and outdoor flies. The indoor flies were fed food that was made with yeast as an ingredient, whereas the outdoor cages received yeast from an initial inoculation and then relied on natural growth. We demonstrated that supplementing lab-reared flies with live yeast resulted in a slightly decreased cold hardening response (**Figure 5B; Table 3**). In contrast, previous work found that supplementation with live yeast increases basal (not precooled) cold tolerance (Colinet and Renault 2014). Our results therefore suggest that dietary yeast may have a different influence on basal cold tolerance as opposed to cold hardened cold tolerance.

The role of yeast in cold hardening may be linked to its role in modulating life history tradeoffs. Higher yeast availability correlates with lower starvation tolerance, reduced lifespan, and higher fecundity, suggesting that yeast modulates a tradeoff between somatic maintenance and reproduction (Simmons and Bradley 1997; Tu and Tatar 2003; Chippindale *et al* 2004). Taken together, we suggest that feeding flies increased yeast may prompt them to prioritize reproduction over survival, thereby reducing energetic investment in processes related to cold hardening. If the fruit-fed cages had high levels of yeast growth in the summer that diminished in the late fall, these changes could have contributed to seasonal variation in cold hardening. Our observation that dietary yeast influences the cold hardening response is further evidence that cold hardening is a plastic phenotype that responds to nutritional conditions.

#### Effects of life history traits on the cold hardening response

The outdoor and indoor cages likely varied in density and age structure, and these factors could plausibly contribute to the observed differences in the cold hardening response. We found that the cold hardening response declines with age in lab-reared flies, perhaps suggesting an age-dependent mechanism (**Figure 5C**). Cold hardening occurs in larvae, pupae, and adult flies, but adults appear to exhibit the greatest cold hardening ability (Czajka and Lee 1990). Previous studies have demonstrated that increased age correlates with increased chill coma recovery time and decreased cold tolerance (David *et al* 1998; Colinet et al 2013). Taken together, these data suggest that the ability to cold harden increases over the course of the fly’s development and eventually tapers off in late adulthood as a result of age-related decline. Without knowing the specifics of age structure in the outdoor cages, it is difficult to conclude how age may have influenced cold hardening in the field-collected samples. However, our laboratory data and the work of others suggest that it may have been a factor, and thus aging serves as another example of the plasticity of cold hardening.

The outdoor cages contained a large volume of fruit, perhaps resulting in lower larval densities relative to the indoor controls. We found that high developmental density results in an increased cold hardening response (**Figure 5D**). Previous work has shown that high larval density induces increased heat tolerance (Sorenson and Loeschcke 2001) and cold tolerance (Henry *et al* 2018). Notably, larval crowding has also been shown to increase adult fat content (Zwaan *et al* 1991) which, as discussed above, may result in greater cold hardening ability. Therefore, the likely higher densities experienced by cornmeal-molasses-fed flies may have primed them for improved cold hardening relative to fruit-fed flies that likely experienced lower densities. The impact of larval density on cold hardening, combined with the influences of age and nutrition, point to the highly plastic nature of this trait.

## Conclusions

Short-lived organisms in changing environments face two options for survival: plastic physiological responses or adaptive tracking. While *D. melanogaster* exhibits seasonal adaptive tracking for several phenotypes (Schmidt and Conde 2006; Behrman *et al* 2015, 2018), we found no evidence for genetic changes in cold hardening in flies experiencing natural seasonal conditions. Instead, cold hardening is highly dependent on a variety of environmental and life history conditions. Understanding the use of plasticity versus adaptive tracking is critical for modelling and predicting how organisms will cope with a changing climate and the associated shifts in environment and habitat range (Hoffmann and Sgrò 2011; Merila and Hendry 2014; Stoks *et al* 2014; Oostra *et al* 2018). Based on our data, we concur with Ayrinhac *et al* (2004) that the factors that influence plasticity may be more important than standing genetic variation for some organisms facing thermal extremes.

## Acknowledgements

The authors would like to thank Dr. Sarah Kucenas, Dr. Laura Galloway, and Dr. Amanda Gibson for their input on drafts of this manuscript. We thank Daniel Song for assistance with field work. We thank members of the Bergland Lab for their comments and assistance. This work was funded by award #61-1673 from the Jane Coffin Childs Memorial Fund for Medical Research (to PAE), an NIH NIGMS grant (R35 GM 119686 to AOB), a Harrison Undergraduate Research Award from the University of Virginia (to HMS), and start-up funds provided by the University of Virginia (to AOB).

## Data Accessibility Statement

All data and R scripts used for analysis and plotting will be made available on Data Dryad.

## Author Contributions

AOB, PAE, and HMS generated the experimental design. HMS and PAE carried out experiments. PAE and HMS conducted analysis and summaries of the results. HMS wrote the first draft of the manuscript and HMS, PAE, and AOB edited and finalized the manuscript. HMS, PAE, and AOB procured funding.

